# Repeat aware evaluation of scaffolding tools

**DOI:** 10.1101/148932

**Authors:** Igor Mandric, Sergey Knyazev, Alex Zelikovsky

## Abstract

**Summary:** Genomic sequences are assembled into a variable, but large number of contigs that should be scaffolded (ordered and oriented) for facilitating comparative or functional analysis. Finding scaffolding is computationally challenging due to misassemblies, inconsistent coverage across the genome, and long repeats. An accurate assessment of scaffolding tools should take into account multiple locations of the same contig on the reference scaffolding rather than matching a repeat to a single best location. This makes mapping of inferred scaffoldings onto the reference a computationally challenging problem. This paper formulates the repeat-aware scaffolding evaluation problem which is to find a mapping of the inferred scaffolding onto the reference maximizing number of correct links and proposes a scalable algorithm capable of handling large whole-genome datasets. Our novel scaffolding validation pipeline has been applied to assess the most of state-of-the-art scaffolding tools on the representative subset of GAGE datasets.

**Availability:** The source code of this evaluation framework is available at https://github.com/mandricigor/repeat-aware. The documentation is hosted at https://mandricigor.github.io/repeat-aware.

## 1 Introduction

Genome sequencing is a vital component for detailed molecular analysis of any organism. Although the depth of sequence coverage grows, the genomic sequence is still assembled into a variable, but large number of contigs. Correct inference of the order and orientation of contigs (referred to as a scaffolding problem) greatly facilitate analysis of gene order and synteny, comparative or functional genomics or investigating patterns of recombination. Interruptions in contig assembly are caused by sequencing technology drawbacks (coverage variation, sequencing bias, and limited read lengths) and the existence of repeats in the genome. While advances in sequencing technologies can reduce technological drawbacks, the long repeats will still be a challenge in foreseeable future.

Genome assembly is one of the oldest, yet still one of the most relevant problems in bioinformatics even nowadays. Traditionally, any genome assembly pipeline is divided into three stages: contig assembly, scaffold assembly, and gap filling. Scaffold assembly is the problem of building chains of contigs from the information provided by paired-end reads. Since early 2000s many scaffolding problem formulations and algorithms were proposed: OPERA [7], SSPACE [4], BESST [17], ScaffMatch [16]. Most of the scaffolding formulations imply the construction of so-called scaffolding graph *G* = (*V*, *E*), where *V* is usually the set of contigs (or contig strands [16]) and E is the set of links obtained by grouping multiple paired-end reads aligning to different contigs. Some scaffolding tools provide heuristics for building paths in *G* (SSPACE, BESST), where each path corresponds to a scaffold. Optimization approaches reduce scaffolding to maximizing the number of correct links or minimizing the number of erroneous links. Such formulations are usually NP-hard [7, 16].

Finding true scaffolding is computationally challenging due to different factors: mis-assemblies, inconsistent coverage across the genome, but the most important challenge though is the presence of genome repeats. For example, the human genome is reported to contain up to 50% of repeated sequences [20]. Contig assembly tools are not able to distinguish different copies of the same repeat, therefore, all the copies of the same repeated DNA region are usually collapsed into one contig. This creates multiple erroneous links in the scaffolding graph. On the other hand, the reference scaffolding should split such contig into several copies in order to correctly correspond to the reference genome. Therefore, an accurate evaluation of an inferred scaffolding should take into account multiple locations of the same contig on the reference scaffolding rather than matching a repeat to a single best location. This makes mapping of an inferred scaffolding onto the reference scaffolding a nontrivial problem.

There are numerous slightly conflicting parameters for evaluating scaffolding quality – N50, corrected (or true) N50, number of correct and incorrect links between contigs with corresponding sensitivity and PPV, number of inverted contigs, number of correctly assembled genes, etc. (see e.g., [8, 16, 12, 2, 15]). Notoriously, the most difficult to maximize is true N50, which is the minimum length of inferred scaffolds which cover 50% of the genome. Here we focus on a simpler objective of maximizing the number of correctly inferred links between contigs.

### The Scaffolding Evaluation Problem

Find a mapping of the inferred scaffolding onto the reference maximizing number of correct contig links (contig connections, joins).

The formal definition of scaffolding link will be given in Section 3.

Let us view contigs as genes, repeats as gene families, inferred scaffolds and reference scaffoldings as two genomes over the same set of gene families. Then the Scaffolding Evaluation (SE) problem is equivalent to finding the breakpoint distance [ 19, 1] between two signed sequences corresponding to the inferred and reference scaffoldings (see Figure 1). Computing the breakpoint distance is NP-hard [3] in presence of nontrivial gene families and, therefore, the SE problem is also NP-hard in presence of repeated contigs.

**Figure 1:**
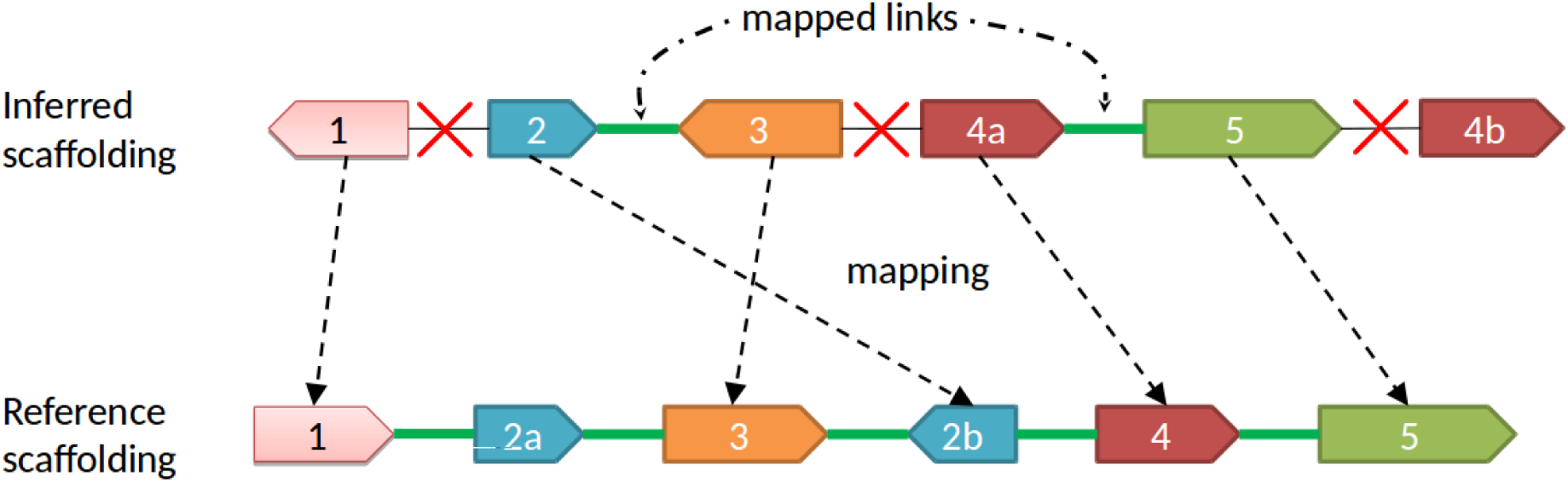
Matching of contigs in the scaffolding S and in the reference *R*. Only the links (2,3) and (4a,5) are mapped correctly. Contig 2 has two copies 2a and 2b in the reference scaffolding, contig 4 is inferred to have two copies 4a and 4b. Assigning contig 2 to either of the copy 2a or 2b, as well as assigning either 4a or 4b to the reference contig 4 affects the number of correctly inferred contigs links. Indeed, assigning contig 2 to the reference contig 2a and contig 4b to 4 will mistakenly undercount the number of correct links. The optimal assignment (2 to 2b, 4a to 4) allows to detect two correctly linked contig pairs.

The rest of the paper is organized as follows: Section 2 provides background and motivation for a new scaffolding validation framework, in Section 3 we give Integer Linear Program based formulation of the Scaffolding Evaluation Problem. Sections 4.1-4.3 explain the validation pipeline and provide the comparison results of most state-of-the-art scaffolding tools.

## 2 Background

The traditional way of measuring the quality of a scaffold assembly most widely used by practitioners used to be N50 which is the shortest sequence length at 50% of the genome. Although this measure is intuitively clear and allows one to estimate the contiguity of the scaffolds, it is rather meaningless: all the contigs randomly joined together produce a scaffolding with highest N50. An alternative way to measure contiguity of scaffolds is to apply corrected N50 metric which is N50 computed after removing all the wrong contig connections.

One of the most recent frameworks for scaffolding evaluation [8] proposed a strategy for evaluating the output of scaffolding tools based on a known ground truth. In [8], the so-called “assembly” contigs (i.e., produced by an assembly tool, for example, Velvet [21]) used to produce “perfect” contigs. The procedure for obtaining the set of perfect contigs is to align the “assembly” contigs to the reference genome with nucmer [5], merge all overlapping hits corresponding to one contig and removing contigs which are completely contained inside any other contig. Then, from each perfect contig a unique sequence tag is extracted as anchor for identification of mutual ordering, orientation, and distance of contigs in the output of scaffolding by aligning the tags to them. In [8], the perfect contigs were not assumed to be repeated ones or to have a sub-part which is repeated in the genome. The only best nucmer hit of each assembly contig was considered. Evaluation [8] did not take into account repeated contigs, though a solid evidence for a high number of repeats exists. For example, in the benchmarking contig datasets which we will use in this paper (see Table 1) percentage of repeated contigs is considerable. Thus, the percentage of repeated contigs in the *S. aureus* dataset is 18.4%.

**Table 1:** The number of unique contigs and the number of repeats in three GAGE datasets.

|  | Unique contigs | Contigs with 2 copies | Contigs with 3 copies | Contigs with 4 copies | Contigs with 5 copies | Total number of contigs |
| --- | --- | --- | --- | --- | --- | --- |
| *S. aureus* | 164 | 4 | 2 | 2 | 3 | 201 |
| *R. sphaeroides* | 572 | 21 | 4 | 2 | 0 | 634 |
| *H. sapiens (chr 14)* | 44748 | 533 | 37 | 8 | 5 | 45982 |

Taking into consideration only the best hit creates the possibility of mis-calling the right link between two contigs in a scaffolding output dataset. Consider the following example. We ran SSPACE on the S. aureus dataset from the GAGE [18] project. All the contigs in the ground truth answer were enumerated from 1 to 170 following framework [8]. In the scaffolding, SSPACE placed the contig 102 (463 bp long) between contigs 19 (57049 bp long) and 20 (132320 bp long) and this situation was classified as producing two errors, namely, incorrect orientation and incorrect distance (see Figure 2). If we re-align contig 102 to the S. aureus genome using nucmer with the parameter of similarity score set to 97%, we notice that contig 102 has three potential placements: a) between contigs 19 and 20 with similarity score 98.49% and it is reversed, b) between contigs 79 and 80 with similarity score 99.78%, and c) between contigs 101 and 103 with similarity score 100% - the actual “best hit” placement chosen as a perfect contig in [8]. Thus, contig 102 in fact has 3 copies under the similarity level of 97%, and the decision taken by SSPACE to place 102 between 19 and 20 is “not that wrong”. Exactly the same mis-classification of the contig links in SSPACE output is done with contig 100 which is placed between contigs 22 and 23 by SSPACE. In fact, under the 97% threshold, one can observe 5 copies: a) between 22 and 23 with similarity 98.23%, b) between 94 and 95 with similarity 99.05%, c) between 97 and 98 with similarity 98.10%, d) between 99 and 101 with similarity 100%, and e) between 130 and 131 with similarity 98.10%. Again, contig 100 added two more errors to the evaluation.

**Figure 2:**
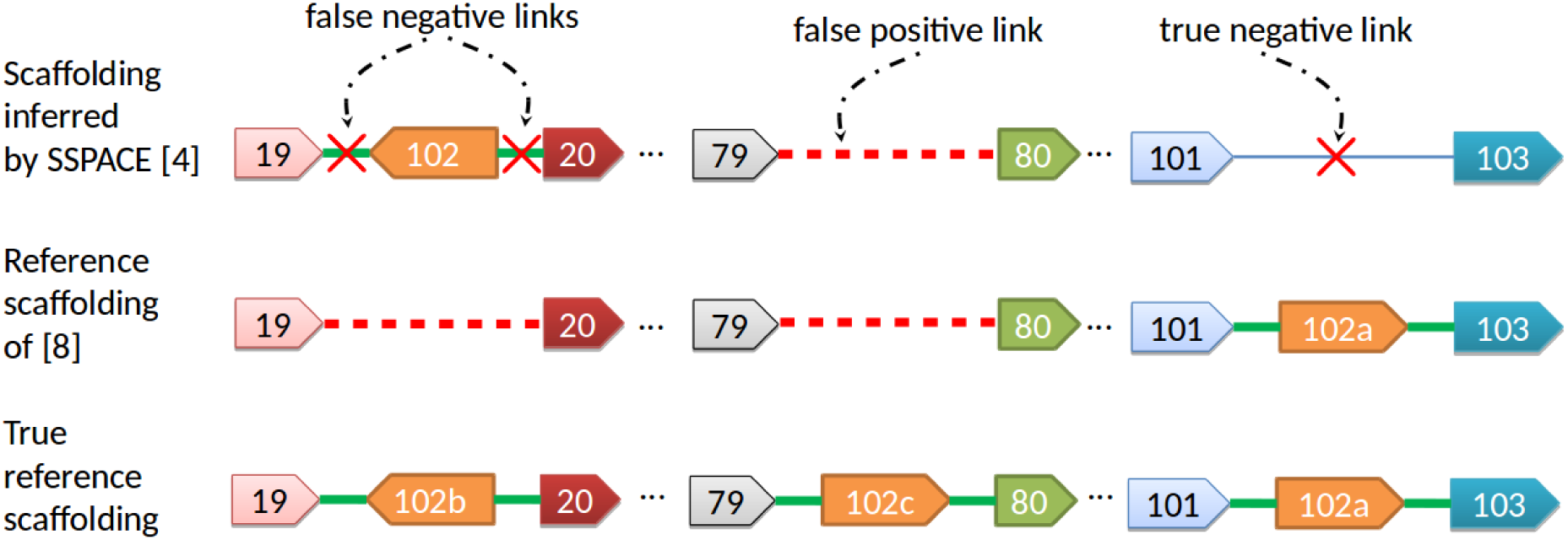
The reference scaffolding contains three copies of contig 102 (namely, 102a, 102b, and 102c). In the reference scaffolding of Hunt et al. [8] only the best hit is considered and two copies with a high identity level (> 97%) are discarded. As a result the contig 102 is treated as erroneously placed by SSPACE between contigs 19 and 20 resulting in two false negative links. Similarly, since the contig 102c is missing, the link between contigs 79 and 80 is falsely treated as correct. Finally, the missing contig 102a is correctly classified.

Another problem with evaluation [8] appears when there are repeated sequences in two or more different chromosomes. Repeated contigs may be “shared” between different genomic sequences.

Although each contig, as is shown above, may have multiple copy numbers in genomes, most of the state-of-the-art scaffolding tools treat contigs produced by assembly tools as having only a single copy. Thus, in order to correctly evaluate scaffolding, one needs to consider all the possible placements of each repeated contig and do not report an error in case when a different contig copy is selected within the defined level of tolerance (we use 97%) rather than the best hit (100%).

As far as we are aware of, OPERA-LG [6] is the only scaffolding tool which handles repeated contigs, i.e. it outputs scaffolds where some of the contigs have multiple copies. As for each contig its copy number in the scaffolding and in the reference may differ, an additional challenge is emerging for evaluating the scaffolding. In [6], the performance of repeat aware scaffolding was assessed by the ability of the scaffolding tool to correctly infer the sequence for gaps where OPERA-LG placed repeats.

In this paper we propose a novel scaffolding evaluation framework which presents a unified approach for assessment of scaffoldings with and without repeated contigs. Our framework as compared to [8] provides an only quantitative measure (number of correct contig links) which is an advantage for an easier comparison of different tools performances. For a more detailed analysis of scaffolding results, one can use more other metrics/scores (see Section 4.4).

## 3 Exact scaffolding evaluation

In this section we formulate Scaffolding Evaluation Problem as an Integer Linear Program (ILP). We start with the formal definitions.

### Definition

Let *C* = {*c*_1_, *c*_2_, …, *c_n_*} be a set of contigs. Sequence 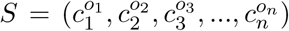, where *o_i_* ∊ {+, −}, is called *scaffold*. Note that *S* may contain multiple copies of any of 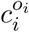 or 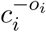 (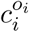 with the opposite orientation). We will call *link* an ordered pair of adjacent contigs 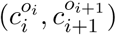. The same link may be alternatively denoted as 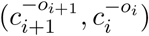 (i.e, these two links are identical, 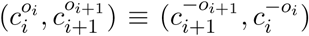. The set of links belonging to a scaffolding S will be denoted as *L_S_*.

*Scaffolding* denotes the set of scaffolds produced by a scaffolding tool. Consider a genome of a model organism (or any other organism for which a reliable reference exists). Let *S* be a scaffolding produced by a scaffolding tool over a set of contigs *C* and let *R* be a scaffolding over the same set of contigs *C* for which the following properties are satisfied:

(i) Any contig 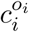 from any scaffold belonging to *R* is encountered in *R* as many times as many non-overlapping alignments of 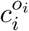 to the genome reference exist within the predefined level of sequence identity;
(ii) All the alignments of all the contigs to the genome reference are consistent with *R*.

For convenience, we will refer to *R* as simply *reference R*. Ideally, the objective of any scaffolding tool is to output a scaffolding *S* which will be identical with *R*, or at least “as close as possible” to *R*. Intuitively, this means that one wants to minimize the number of erroneous links in S when being compared to R, or to maximize the number of valid links. Link 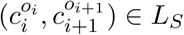 is valid if either 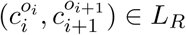 or 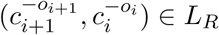.

We formulate the following ILP:
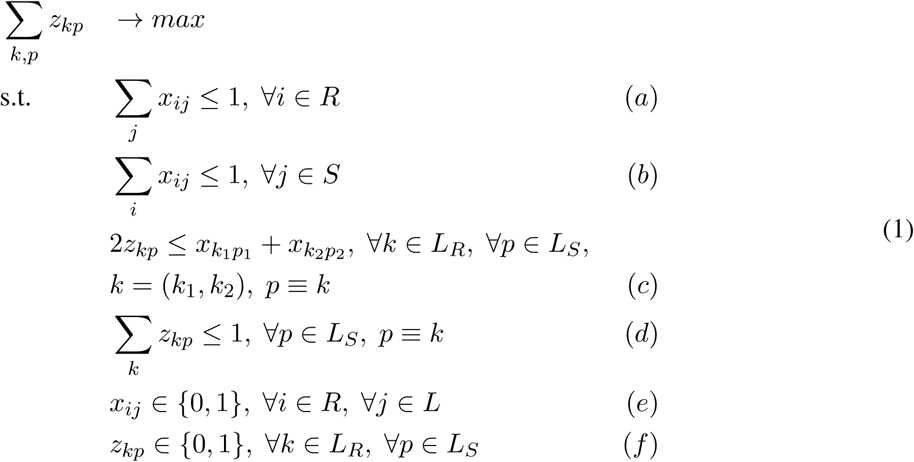

Binary variables *x_ij_* ∊ {0, 1} have the following meaning:

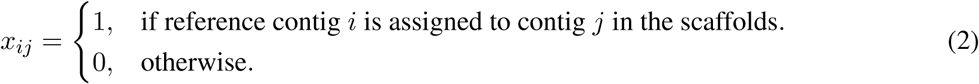

Conditions (a) and (b) from the ILP (1) guarantee that each contig *j* ∊ *S* is assigned to at most one contig i ∊ *R*.

For any link *k* = (*k*_1_, *k*_2_) ∊ *L_R_* and for any link *p* ∊ *L_S_* which is identical with *k* we introduce a binary variable *z_kp_* ∊ {0,1} with the following meaning:

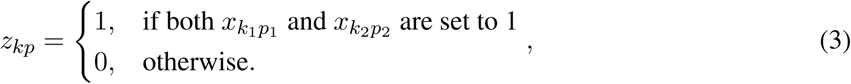

Constraints (c) correspond to the relations between the variables 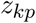 and the variables 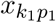 and 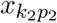. It may happen that for a given link *k* ∊ *R* there may be multiple links *p*_1_, *p*_2_, …, *p_n_* ∊ *S* identical with *k*. To guarantee that at most one link *p_i_* from the scaffolding *S* is matched with *k*, constraint (d) is introduced to the problem.

Maximizing the objective 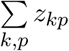 means to find an assignment of scaffold contigs to the reference contigs, i.e. assignment of variables *x_ij_* such that to maximize the number of contig links *k* ∊ *R* matched with identical to them contig links from *S*.

## 4 Repeat-aware validation of scaffolders

### 4.1 Datasets with repeats

As in [8], we use the assembly contigs produced by the Velvet assembler, as the starting point. Instead of identifying the best nucmer hit for each assembly contig, we find all the hits with the minimal identity of 97%. Thus, each assembly contig may have multiple hits. If a contig is fully aligned with the reference genome, we add it to the benchmark contig dataset. We do not take into account smaller parts of assembly contigs which may align to the reference to not to produce too fragmented benchmark contigs. Otherwise, for each reference sequence, we aggregate overlapping nucmer hits coming even from different contigs into one artificial contig and add it to the benchmark contig dataset. We do so because two assembly contig sequences may differ due to assembly errors, but share a significant portion of each other when being aligned to the reference. We “correct” the assembler errors and join these contigs into one. The procedure of building artificial contigs is depicted in Figure 3.

**Figure 3:**
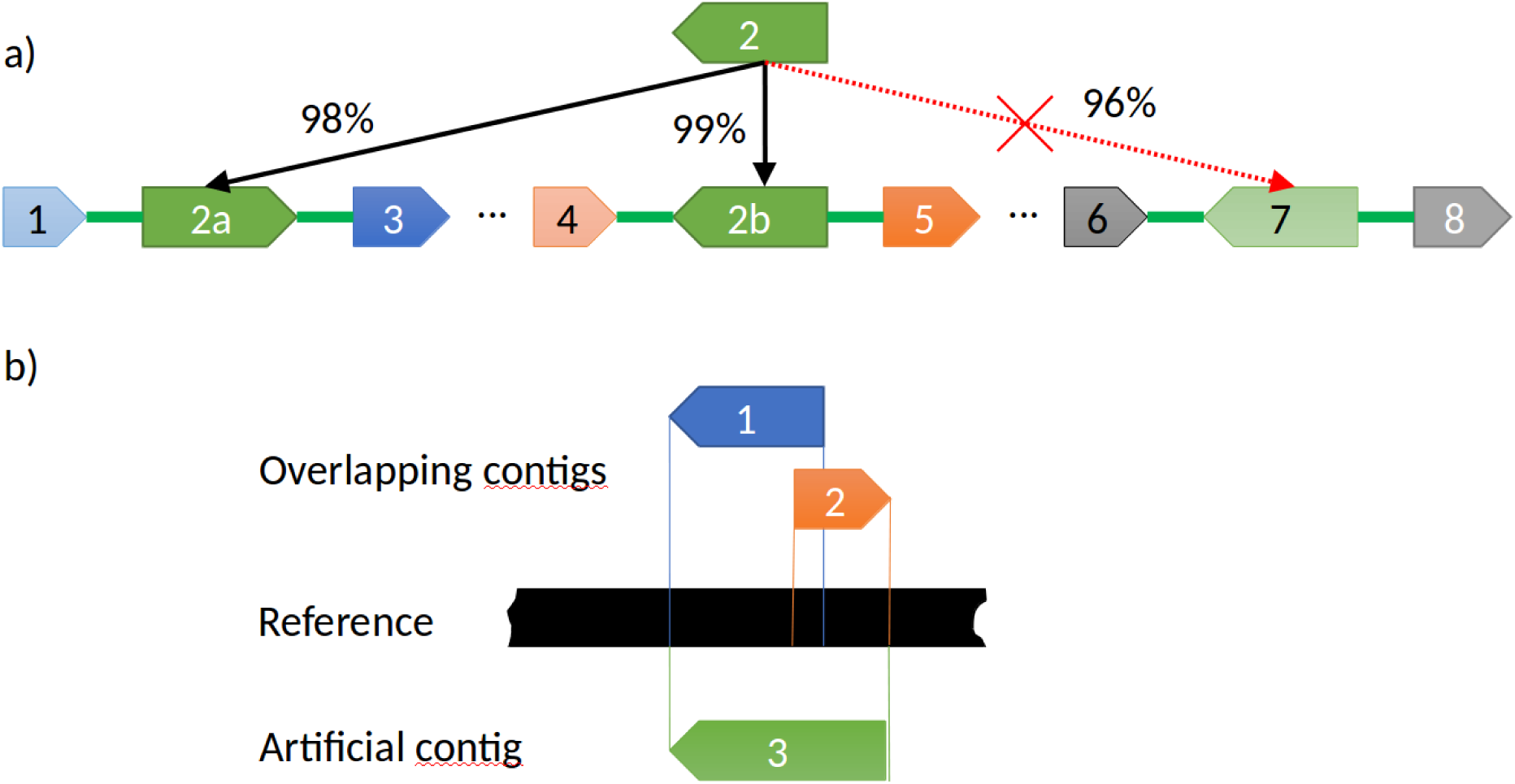
a) Choosing the longest hit of each contig within the predefined level of sequence identity. Contig 2 aligns to three different places on the reference: 2a, 2b, 7. Artificial contig 7 is discarded because of its identity level below the predefined threshold of 97% and its length is less than the length of artificial contigs 2a and 2b. b) Assembly contigs 1 and 2 significantly overlap with each other. We merge such contigs into one artificial contig.

We built three sets of artificial contigs based on Velvet assembly from the GAGE project: *S. aureus*, *R. sphaeroides*, *H. sapiens (chr 14)*. In Table 1 copy number distribution and the total number of contigs for each dataset is presented.

### 4.2 Evaluation pipeline

We implemented our evaluation framework using Python programming language (version 2.7). We used nucmer version 3.1 for aligning contigs to the reference. Integer Linear Program (1) is solved with the aid of IBM CPLEX solver version 12.7. In order to evaluate a scaffolding S given the reference genome, we perform the following sequence of steps:

1. Take assembly contigs and produce perfect contigs as described in Section 4.
2. Build the reference scaffolding *R*
3. Run a scaffolding tool on the perfect contigs and a paired-end read dataset
4. Align the perfect contigs to the output scaffolding using nucmer (with exactly the same identity level which was used for building the artificial contigs in step 1).
5. Build the output scaffolding *S*
6. Solve the Scaffolding Evaluation Problem with input data (*S*, *R*).

### 4.3 Validated Scaffolding tools

We ran the following scaffolding tools: OPERA-LG (version 2.0.6), OPERA (version 1.4), ScaffMatch (version 0.9), SOAPdeNovo2 [14] (version 2.04), BESST (version 2.2.5), BOSS [13] (latest version from GitHub), SSPACE (version 3.0) on the three artificial contig datasets: *S. aureus*, *R. sphaeroides*, *H. sapiens (chr 14)* from the GAGE project. The following Illumina paired-end read datasets were used: *S. aureus* - read length 37, insert size 3600; *R. sphaeroides* - read length 101, insert size 3700; *H. sapiens (chr 14)* - read length 101, insert size 2700. Most of the tools accept a user specified read aligner (for example, BWA [11], Bowtie [10] or Bowtie2 [9]. As OPERA-LG is bundled and BESST is better to be used with BWA (as per BESST documentation, https://github.com/ksahlin/BESST), we used it for most of the experiments. ScaffMatch was run with Bowtie2 alignments 1).

### 4.4 Validation results

We apply our ILP (1) to obtain the number of inferred correct contig links by each scaffolder. For the largest data set H. sapiens (chr14) the ILP was scalable enough taking less than 15 min on 2.5GHz 16-core AMD Opteron 6380 processors with 256Gb RAM running under Ubuntu 16.04 LTS. In the Table 2 we present the number of correct contig links as well as sensitivity and PPV for the scaffolds produced by each of the tools. Here sensitivity and PPV are computed using the following formulas:

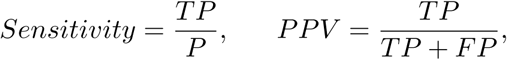

where *P* is the total number of contig links in the reference scaffolding with repeats which is the number of contigs minus the number of chromosomes, *TP* is the number of correct contig links in the scaffolding output (true positives) and *FP* is the number of inferred erroneous links (false positives).

**Table 2:**
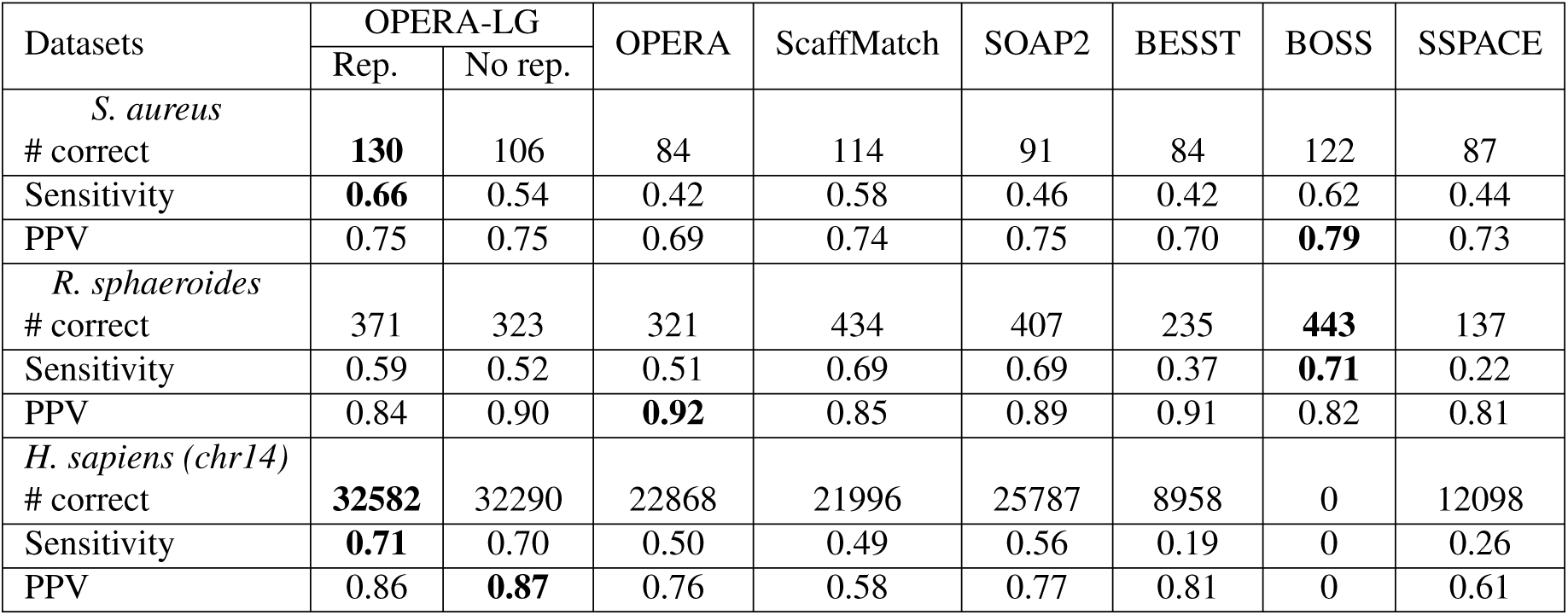
Number of correct contig links, sensitivity and positive predictive value (PPV) on GAGE datasets as obtained from solving the ILP (1)

| Datasets | OPERA-LG |  | OPERA | ScaffMatch | SOAP2 | BESST | BOSS | SSPACE |
| --- | --- | --- | --- | --- | --- | --- | --- | --- |
|  | Rep. | No rep. |  |  |  |  |  |  |
| *S. aureus* |  |  |  |  |  |  |  |  |
| # correct | **130** | 106 | 84 | 114 | 91 | 84 | 122 | 87 |
| Sensitivity | **0.66** | 0.54 | 0.42 | 0.58 | 0.46 | 0.42 | 0.62 | 0.44 |
| PPV | 0.75 | 0.75 | 0.69 | 0.74 | 0.75 | 0.70 | **0.79** | 0.73 |
| *R. sphaeroides* |  |  |  |  |  |  |  |  |
| # correct | 371 | 323 | 321 | 434 | 407 | 235 | **443** | 137 |
| Sensitivity | 0.59 | 0.52 | 0.51 | 0.69 | 0.69 | 0.37 | **0.71** | 0.22 |
| PPV | 0.84 | 0.90 | **0.92** | 0.85 | 0.89 | 0.91 | 0.82 | 0.81 |
| *H. sapiens (chr14)* |  |  |  |  |  |  |  |  |
| # correct | **32582** | 32290 | 22868 | 21996 | 25787 | 8958 | 0 | 12098 |
| Sensitivity | **0.71** | 0.70 | 0.50 | 0.49 | 0.56 | 0.19 | 0 | 0.26 |
| PPV | 0.86 | **0.87** | 0.76 | 0.58 | 0.77 | 0.81 | 0 | 0.61 |

Also, in Table 3 we present results for corrected N50 which is a widely used state-of-the-art metric for scaffold contiguity assessment. Corrected N50 is obtained by breaking the incorrect links and computing N50 metric on the resulting scaffolds.

**Table 3:** Contiguity characteristics of the scaffolding outputs: corrected N50 (K bp) and corrected chain N50; scaffolding chain characteristics - maximum and average chain length in scaffolding outputs.

| Datasets | OPERA-LG |  | OPERA | ScaffMatch | SOAP2 | BESST | BOSS | SSPACE |
| --- | --- | --- | --- | --- | --- | --- | --- | --- |
|  | Rep. | No rep. |  |  |  |  |  |  |
| *S. aureus* |  |  |  |  |  |  |  |  |
| Scaff. corrected N50 (K bp) | 586 | 588 | 1087 | 460 | NA* | 304 | 729 | 262 |
| Corrected chain N50 | **9** | 5 | 3 | 7 | 3 | 3 | 8 | 3 |
| Maximum chain length | **29** | 25 | 19 | 24 | 12 | 11 | 27 | 24 |
| Average chain length | 3.28 | 2.55 | 1.93 | 2.90 | 2.26 | 1.93 | **3.39** | 2.00 |
| *R. sphaeroides* |  |  |  |  |  |  |  |  |
| Scaff. corrected N50 (K bp) | 194 | NA* | NA* | 2532 | 2522 | 44 | 484 | 27 |
| Corrected chain N50 | 7 | 5 | 5 | **37** | 18 | 2 | 23 | 1 |
| Maximum chain length | 41 | 37 | 37 | **238** | 236 | 28 | 123 | 7 |
| Average chain length | 2.60 | 2.17 | 2.16 | 3.66 | 3.36 | 1.65 | **3.85** | 1.30 |
| *H. sapiens (chr14)* |  |  |  |  |  |  |  |  |
| Scaf. corrected N50 (K bp) | 64 | 66 | 82 | 17 | 85 | 2 | NA** | 9 |
| Corrected chain N50 | **12** | 11 | 3 | 3 | 6 | 1 | NA** | 1 |
| Maximum chain length | **107** | **107** | 38 | 67 | 48 | 53 | NA** | 27 |
| Average chain length | **3.56** | 3.48 | 2.02 | 1.98 | 3.12 | 1.24 | NA** | 1.36 |

\* Program crashed: cause unclear.
\*\* Scaffolder produced no contig links.

We introduce two metrics based on the length of scaffolded contig chains: *average corrected chain length* and *corrected chain N50*. Average corrected chain length is computed by breaking all the incorrect contig links and computing the average chain length of the resulting scaffolds. Chain length is equal to the number of its contigs. Similarly, corrected chain N50 is computed as corrected N50 of the chain lengths. The results are reported in Table 3.

The results display a very broad range of correctly assembled links. OPERA-LG with enabled repeated scaffolding (the option *filter*_*repeat* = *no*) (OPERA-LG/Rep.) finds exceedingly more correct contig links than all other tools including OPERA-LG with the option *filter*_*repeat* = *yes* denoted OPERA-LG/No rep. On the other hand, PPV of OPERA-LG/Rep. is worse than OPERA-LG/No rep. because it sometimes adds too many copies of repeats. Unfortunately, BOSS did not connect contigs for the human dataset.

### 4.5 Classification of incorrect links

Since our framework is targeted at optimizing the number of correct links, it is of particular interest for genome assembly practitioners to be able to classify wrong links in the output of any scaffolding tool. Knowledge of incorrect types of links may provide an additional insight into drawbacks of scaffolding algorithms.

In [8] framework, the following types of incorrect links were distinguished:

1. Two contigs originate from same reference sequence, but their orientation in the scaffolds is incorrect;
2. Two contigs originate from different reference sequences;
3. Two contigs originate from the same reference sequence but are the wrong distance apart;
4. Two contigs originate from the same reference sequence but are not in the correct order.
5. Pair of contigs that originated from the same reference sequence, but their orientation and order were incorrect;
6. Two contigs were from the same reference sequence, but were the wrong distance apart and in the wrong order.

In our classification we distinguish the following cases:

- Wrong copy link (C) - a contig link incident to at least one wrong copy of a contig (see link *D* in the Figure 4);
- Different references (R) - a contig link connecting contigs coming from different reference sequences. It is equivalent to 1) from above classification.
- Wrong order/orientation (ORD-ORI) - the link is not equivalent to the correct link in terms of ordering/orientation of its contigs (see, for example, link *B* in Figure 4). This case is equivalent to 1) and/or 4) and/or 5) from the above classification. We do not distinguish between these three cases. Consider a contig link (*A*^+^, *B^−^*) (same as (*B*^+^, *A^−^*)). Then, an incorrect link (*B^−^*, *A*^+^) may be classified as either a wrong order link (we have to swap *A*^+^ and *B^−^*, or incorrect orientation (we have to switch the orientation of both contigs).
- Jumping link (J) - the two contigs are in correct order and orientation, but at least one contig is missing between them in the reference. This corresponds to the case of “skipped” contigs in [8]. It is treated as a correct link in [8], we do not take such links into account in our ILP model. Note that we include here the case 3) from the above classification.
- Jumping and wrong order/orientation (J + ORD-ORI) - the combination of two cases.

**Figure 4:**
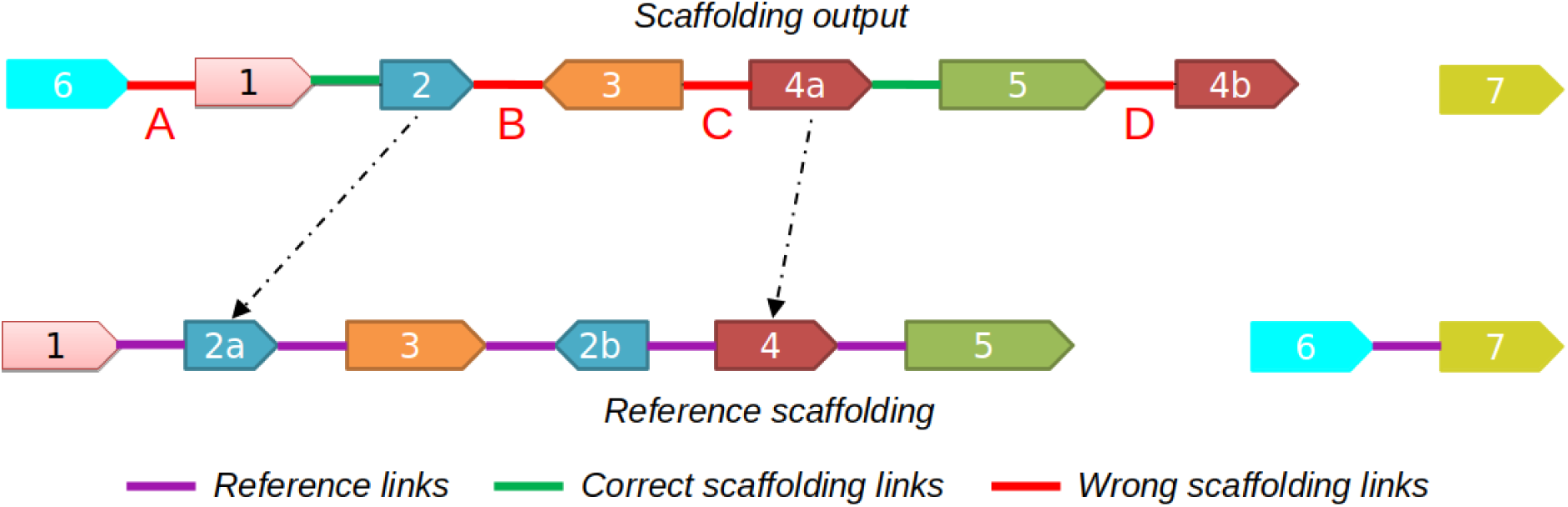
Classification of incorrect links. Contig 2 in the scaffolding output is assigned to contig 2a in the reference, contig 4a in the scaffolding output is assigned to contig 4 in the reference (marked with arrows). There are two correct links (marked with green) and 4 wrong links (marked with red) - *A*, *B*, *C*, *D*. Link A connects contigs 6 and 1 coming from different reference sequences. Link *B* connects contigs 2 and 3 which are not in correct order/orientation. Jumping link *C* connects contigs 3 and 4a which are not in correct order/orientation. Link *D* connects contig 5 with an “extra” copy of contig 4 (namely 4b).

Note that in general, number of R errors in our framework should be less than in [8] since we do admit one contig sequence to be shared among different reference sequences. Also, a new class of wrong links arises in our framework, namely links connecting extra copies of contigs (C error). The results of classification of incorrect links are provided in the Table 4.

**Table 4:** Classification of incorrect links on three GAGE datasets scaffolding outputs: *S. aureus*, *R. sphaeroides*, *H. sapiens (chr14)*. The number of incorrect links of the following types are reported: C - links incident to “extra” contig copies, R - links connecting contigs originating from different reference sequences, J - jumping links (contigs are in correct relative order and orientation, but one or more contigs are skipped in between), ORD-ORI - links connecting contigs in wrong a ordering/orientation, J + ORDORI - a combination of J and ORD-ORI types, namely these are links connecting two contigs in a wrong ordering/orientation and one or more contigs are skipped in between.

| Datasets | OPERA-LG |  | OPERA | Scaffmatch | SOAP2 | BESST | BOSS | SSPACE |
| --- | --- | --- | --- | --- | --- | --- | --- | --- |
|  | Rep. | No rep. |  |  |  |  |  |  |
| *S. aureus* |  |  |  |  |  |  |  |  |
| C | 17 | 0 | 0 | 0 | 0 | 0 | 0 | 0 |
| R | 1 | 0 | 0 | 0 | 0 | 0 | 0 | 0 |
| J | 0 | 0 | 0 | 0 | 0 | 0 | 0 | 0 |
| ORD-ORI | 1 | 1 | 0 | 0 | 1 | 0 | 0 | 0 |
| J + ORD-ORI | 23 | 33 | 36 | 40 | 22 | 36 | 29 | 32 |
| *R. sphaeroides* |  |  |  |  |  |  |  |  |
| C | 34 | 0 | 0 | 0 | 0 | 0 | 0 | 0 |
| R | 1 | 2 | 2 | 6 | 2 | 0 | 25 | 2 |
| J | 0 | 0 | 0 | 0 | 0 | 0 | 0 | 0 |
| ORD-ORI | 0 | 0 | 0 | 4 | 0 | 0 | 7 | 0 |
| J + ORD-ORI | 33 | 31 | 25 | 67 | 32 | 22 | 64 | 31 |
| *H. sapiens (chr14)* |  |  |  |  |  |  |  |  |
| C | 312 | 0 | 0 | 0 | 0 | 0 | N/A | 0 |
| R | 0 | 0 | 0 | 0 | 0 | 0 | N/A | 0 |
| J | 0 | 0 | 0 | 0 | 0 | 0 | N/A | 0 |
| ORD-ORI | 308 | 272 | 10 | 510 | 195 | 11 | N/A | 2 |
| J + ORD-ORI | 4693 | 4672 | 7102 | 15683 | 2481 | 2084 | N/A | 7847 |

Note that in general, number of R errors in our framework should be less than in [8] since we do admit one contig sequence to be shared among different reference sequences. Also, a new class of wrong links arises in our framework, namely links connecting extra copies of contigs (C error). The results of classification of incorrect links are provided in the Table 4.

### 4.6 Comparison of the two evaluation frameworks

In order to demonstrate that our framework allows a more accurate evaluation of scaffolding results, we ran both evaluations on the set of “perfect” scaffoldings used in our experiments. As our framework is aware of repeats, it identifies the perfect scaffoldings as having no erroneous links. Conversely, the Hunt et al. framework does not consider a link with at least one repeated contig placed in an alternative location as a correct one. As a result, it penalizes even the “perfect” scaffoldings treating them as containing multiple incorrect contig joins. In the Table 5 the number of correct and incorrect links inferred by the Hunt et. al framework is reported. For example, on *S. aureus* dataset, the mis-classification rate is almost 9%.

**Table 5:** Errors reported by the Hunt’s framework on the perfect scaffolding datasets

| Dataset | Correct links | Wrong links | Percent wrong |
| --- | --- | --- | --- |
| *S. aureus* | 157 | 15 | 8.7 |
| *R. sphaeroides* | 547 | 42 | 7.1 |
| *H. sapiens* | 44219 | 969 | 2,1 |

## 5 Conclusions

Scaffolding is very important for obtaining qualitative, error free genomes. Not of less importance is validation of scaffolding assemblies. It allows to recognize the strong points of the existing tools and emphasize their drawbacks, thus leveraging future research in the area.

In this paper we presented a novel scaffolding evaluation pipeline which is able to take into account repeated contig sequences. It allows to adequately measure the quality of scaffolding assemblies produced by the current state-of-the-art tools as well as the tools adjusted to handle repeats such as OPERA-LG. We believe that the proposed pipeline will make more attractive developing of repeat-aware releases of existing scaffolding tools.

